# Folding stabilities of ribosome-bound nascent polypeptides probed by mass spectrometry

**DOI:** 10.1101/2022.11.07.515500

**Authors:** Ruiyue Tan, Margaret Hoare, Kevin A. Welle, Kyle Swovick, Jennifer R. Hryhorenko, Sina Ghaemmaghami

## Abstract

The folding of most proteins occurs during the course of their translation while their tRNA-bound C-termini are embedded in the ribosome. How the close proximity of nascent proteins to the ribosome influences their folding thermodynamics remains poorly understood. Here, we have developed a mass spectrometry-based approach for determining the stabilities of nascent polypeptide chains using methionine oxidation as a folding probe. This approach enables quantitative measurements sub-global folding stabilities of ribosome nascent chains (RNCs) within complex protein mixtures and extracts. To validate the methodology, we analyzed the folding thermodynamics of three model proteins (DHFR, CheY and DinB) in soluble and ribosome-bound states. The data indicated that the ribosome can significantly alter the stability of nascent polypeptides. Ribosome-induced stability modulations were highly variable among different folding domains and were dependent on localized charge distributions within nascent polypeptides. The results implicated electrostatic interactions between the ribosome surface and nascent polypeptides as the cause of ribosome-induced stability modulations. The study establishes a robust proteomic methodology for analyzing localized stabilities within ribosome-bound nascent polypeptides and sheds light on how the ribosome influences the thermodynamics of protein folding.

## Introduction

Within a cell, proteins are synthesized as linear polymers of amino acids and subsequently fold into compact three-dimensional structures. The free energy difference between the folded and unfolded states of a protein (Δ_Gfolding_ or thermodynamic folding stability) dictates the fraction of the protein that is in the folded conformation at equilibrium^1^. Folding stability is a physical parameter that is widely divergent between proteins and can range from -20 kcal/mol for stably folded proteins to positive values for natively unstructured proteins^2–4^. Thermodynamic stability is a critical property of proteins, and mutations and environmental conditions that alter folding stabilities have been implicated in a number of protein misfolding diseases^5^.

To date, nearly all analyses of protein folding thermodynamics have been conducted for purified full-length protein *in vitro*. However, the rate of protein folding is generally much faster than the rate of protein translation, and proteins are known to fold during the course of their synthesis as they are being translated by the ribosome^6-9^. During translation, the incompletely synthesized nascent polypeptide chain has the opportunity to explore its conformational landscape in the absence of its entire sequence, while attached by its C-terminus to the ribosome^10, 11^. Accordingly, it has been shown that translational elongation kinetics can influence the folding of nascent proteins and ribosome-bound enzymes can become active prior to completion of translation^12-14^. These and other studies indicate that within cells, the protein folding reaction occurs for partially-synthesized polypeptides during the course of their synthesis under conditions where they are in close proximity to the ribosome. As such, the thermodynamic principles that govern co-translational folding and stability may be fundamentally different than what has been observed for soluble proteins in traditional *in vitro* folding experiments.

How the ribosome influences the energy landscape and folding of nascent proteins has been explored by a number of recent studies^8, 15, 16^. Biophysical and biochemical approaches including NMR spectroscopy, limited proteolysis and single-molecule optical trapping have provided significant insights into the folding thermodynamics of ribosome nascent chains (RNC)^16–21^. One common theme that has emerged from these studies is the ability of the ribosome to destabilize the native structure of nascent polypeptides during the course of translation^16, 20–22^. It is thought that ribosome-induced destabilization during the course of translation may inhibit incompletely synthesized polypeptides from premature folding that may result in the formation of off-pathway intermediates. However, the mechanistic details of this phenomenon and its generality across the translated proteome remains unclear.

In order to fully delineate the principles that govern co-translational folding, it is necessary to evaluate the effect of the ribosome on a large number of structurally diverse proteins. Indeed, whereas our understanding of protein folding thermodynamics in a soluble state has benefitted from decades of research on hundreds of structurally distinct model proteins^4, 23^, the folding of very few proteins has been investigated in ribosome-bound states. This bottleneck is due in part to the limitations and complexities inherent in methodologies used to interrogate the thermodynamics of RNCs and the difficulty of their application to diverse proteins under complex solution conditions.

To enable the analyses of protein stabilities in complex mixtures, a number of recently developed techniques have sought to investigate folding thermodynamics using probes that can be analyzed by mass spectrometry-based proteomics^24, 25^. As examples, hydrogen/deuterium exchange (HX), hydroxyl radical foot printing and limited proteolysis have been coupled with liquid chromatography-tandem mass spectrometry (LC-MS/MS) to provide effective measures of protein structure and folding^26–29^. More recently, we and others have shown that the oxidation kinetics of buried methionine residues, as quantified by LC-MS/MS, can be used to accurately quantify folding stabilities in an approach referred to as Stability of Proteins from Rates of Oxidation (SPROX)^3, 30^. To date, SPROX has been used to analyze the stability of folding domains within soluble proteins and to examine alterations in denaturation patterns induced by ligand binding^3, 31, 32^. Here, we extend the use of SPROX to analyses of folding thermodynamics of RNCs and demonstrate its ability to measure sub-global folding stabilities under native conditions within complex protein mixtures. The results provide a number of new insights into how the ribosome influences the stability of nascent polypeptides and establish SPROX as a generally applicable methodology for analyzing the folding thermodynamics of ribosome-bound proteins.

## Results

### Measurement of protein stabilities by SPROX and MObB

Figure 1A outlines the general principle of SPROX as a method for measurement of protein stabilities^3, 30^. Methionine is a readily oxidizable amino acid and can be converted to methionine sulfoxide by the oxidizing agent hydrogen peroxide (H_2_O_2_)^33^. The oxidation rate of a methionine residue within a protein is contingent on its local structural environment^34–37^. For a methionine that is fully buried in the protein core, unfolding is required for oxidation. Since the kinetics of protein folding are generally faster than oxidation rates, the folding reaction is a pre-equilibrium that modulates the oxidation rate. Thus, buried methionine residues have a slower rate of oxidation within more stable protein regions^37^.

**Figure 1.**
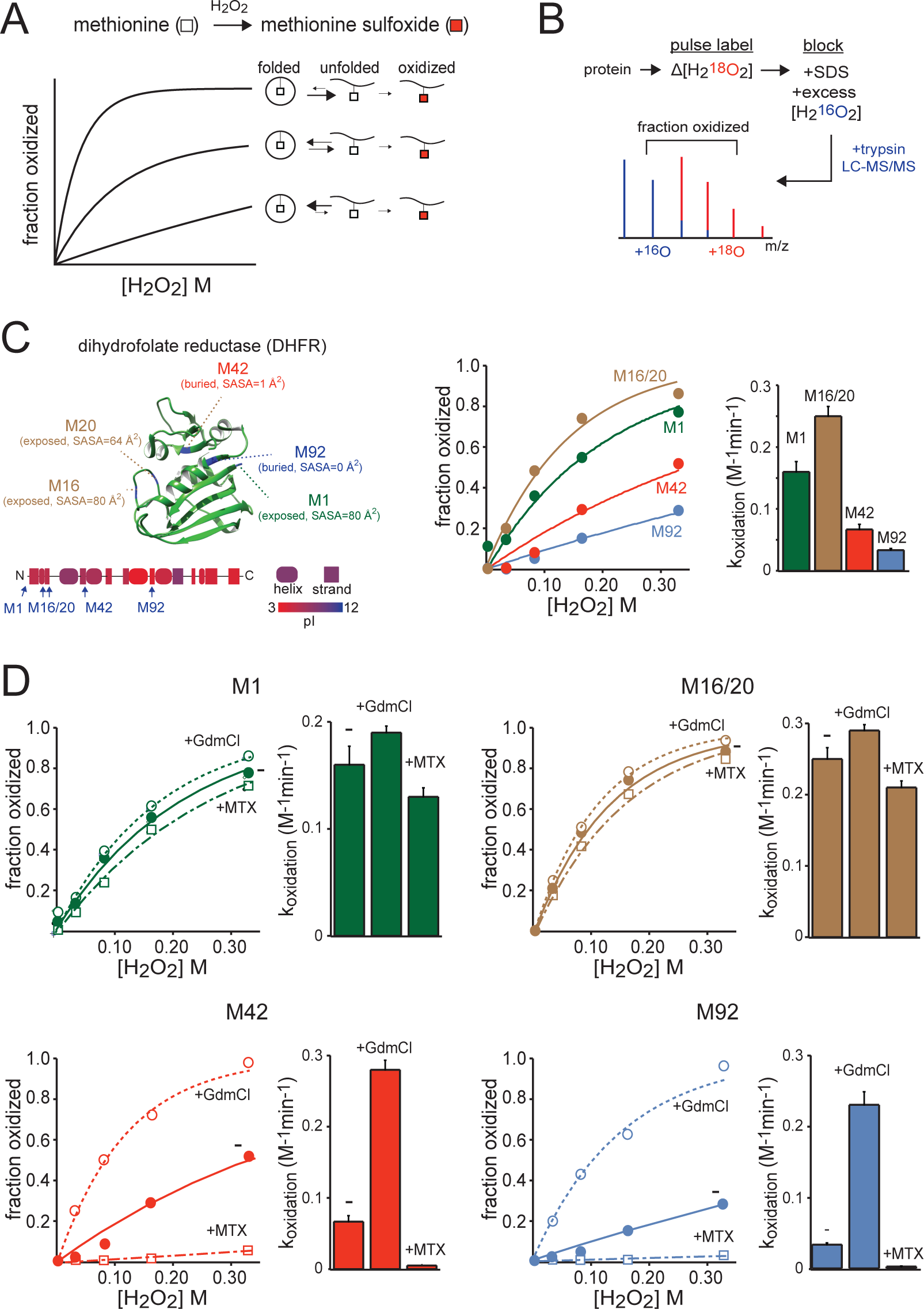
Measurement of protein stabilities by SPROX and validation of the methodology for soluble wildtype DHFR. **A)**Theoretical basis of SPROX. Methionine residues are oxidized by H_2_O_2_ to form methionine sulfoxides. For buried methionine residues, protein unfolding precedes methionine oxidation and therefore protein stabilities modulate oxidation rates. Thus, oxidation rates of buried methionines, determined by quantifying oxidation as a function of [H_2_O_2_], reports on folding stabilities. **B)** Quantitation of methionine oxidation by MObB. Proteins are pulse labeled with varying concentrations of ^18^O H_2_O_2_. After quenching, labeled proteins are denatured, oxidized to saturation with ^16^O H_2_O_2_, digested with trypsin and analyzed by LC-MS/MS. Red and blue lines indicate the spectra of ^18^O- and ^16^O-oxidized peptides, respectively. The ^18^O/^16^O ratio indicates the fractional oxidation of peptides during the pulse phase. **C)** Crystal structure of DHFR (P0ABQ4) highlighting its 5 methionine residues. Solvent accessibility surface area (SASA, Å) indicate that Met 1, 16 and 20 are surface-exposed, whereas Met 42 and 92 are buried in the protein core. Isoelectric point (pI) measurements for each secondary structure element are also illustrated. The plots show measurements of fractional oxidation as a function of [H_2_O_2_] for peptides containing the indicated methionines as well as k_oxidation_ measurements obtained by non-linear fitting of the kinetic data. The error bars indicate the standard errors for parameter estimates based on the fitted model. **D)** Fractional oxidation and k_oxidation_ measurements of DHFR methionines in the presence and absence of MTX (0.5 M) and GdmCl (2.0 M) determined as in (C). The measurements in (C) and (D) were obtained with wildtype soluble full-length DHFR without an AP fusion. See also Figure S1.

Although peptides harboring methionine sulfoxides can be readily detected by mass spectrometry, accurate measurement of fractional methionine oxidation has been historically challenging. This is largely due to the fact that methionine oxidation can spuriously accumulate during the upstream stages of a typical bottom-up proteomics workflow^38^. In previous work, we have developed methodology termed Methionine Oxidation by Blocking (MObB) that circumvents this complication by blocking unoxidized methionines with isotopically labeled oxidants prior to MS analysis^39, 40^. In the experiments described in this study, proteins are initially pulse labeled with ^18^O -labeled H_2_O_2_ and subsequently blocked by exposure to excess levels of ^16^O-labeled H_2_O_2_ prior to MS analysis. Downstream quantitation of the relative ratios of ^18^O and ^16^O oxidized methionine-containing peptides provides an accurate measure of fractional oxidation that occurred during the pulse phase of the experiment (Figure 1B). We used this approach to analyze the oxidation kinetics and folding thermodynamics of a number of model proteins in their soluble and ribosome-bound forms.

### Methionine oxidation kinetics in soluble DHFR

In our initial validation experiments we used *E. coli* dihydrofolate reductase (DHFR) as a model protein. The folding thermodynamics of DHFR can be approximated by a two-state model and has been well studied both in soluble and ribosome-bound forms^21, 41^. We expressed DHFR using the PURE *in vitro* translation system^42^ and analyzed the oxidation kinetics of its methionines by H_2_O_2_ using the strategy outlined above. DHFR contains five methionines: three that are solvent-exposed (M1, M16 and M20), and two that are buried in the protein core (M42 and M92) (Figure 1C). Trypsin digest of DHFR generates MS-detectable peptides that harbor each individual methionine, except for M16 and M20 which are contained in the same tryptic peptide (Figure S1). As expected, the buried methionines M42 and M92 had a significantly faster oxidation rate in comparison to the exposed methionines (Figure 1, S1).

We next repeated the oxidation experiments in the presence two chemicals that are expected to modulate the folding stability of DHFR. Guanidinium hydrochloride (GdmCl) is a chemical denaturant that destabilizes proteins and methotrexate (MTX) is an inhibitor whose binding stabilizes DHFR^43^. As expected, GdmCl and MTX significantly increase and decrease the oxidation rates of the buried methionines M42 and M92 respectively whereas the exposed methionines are unaffected (Figure 1D). Thus, oxidation rates of M42 and M92 are effective indicators of the folding stability of DHFR.

### Methionine oxidation kinetics in ribosome-bound DHFR

To generate RNC complexes of DHFR, we engineered fusion constructs by attachment of the arrest peptide (AP) of *E. coli* SecM protein (Figure 2A). SecM naturally undergoes a strong translational pause resulting from the interaction of the 17-residue AP sequence near its C-terminus with the exit tunnel of the ribosome^44–46^. Fusion of AP to model proteins is a commonly used approach for generating ribosome-bound constructs^16, 18, 19, 21, 47^. We first verified the AP-dependent translational arrest of SecM (Figure S2) and then generated DHFR-AP constructs containing distinct epitope tags at their N- and C-termini (FLAG and HA, respectively). In our initial experiments, DHFR and AP were separated by an unstructured spacer that would be predicted to generate a distance of 50 amino acids between the ribosomal peptidyl transferase center (PTC) and the C-terminus of DHFR following AP-mediated translational arrest (ΔPTC = 50 aa). Assuming that the unstructured spacer has an extended conformation as it traverses the exit tunnel, translationally arrested DHFR in this construct would be expected to protrude from the ribosome and be separated by a sequence of ∼16 residues from the ribosomal surface.

**Figure 2.**
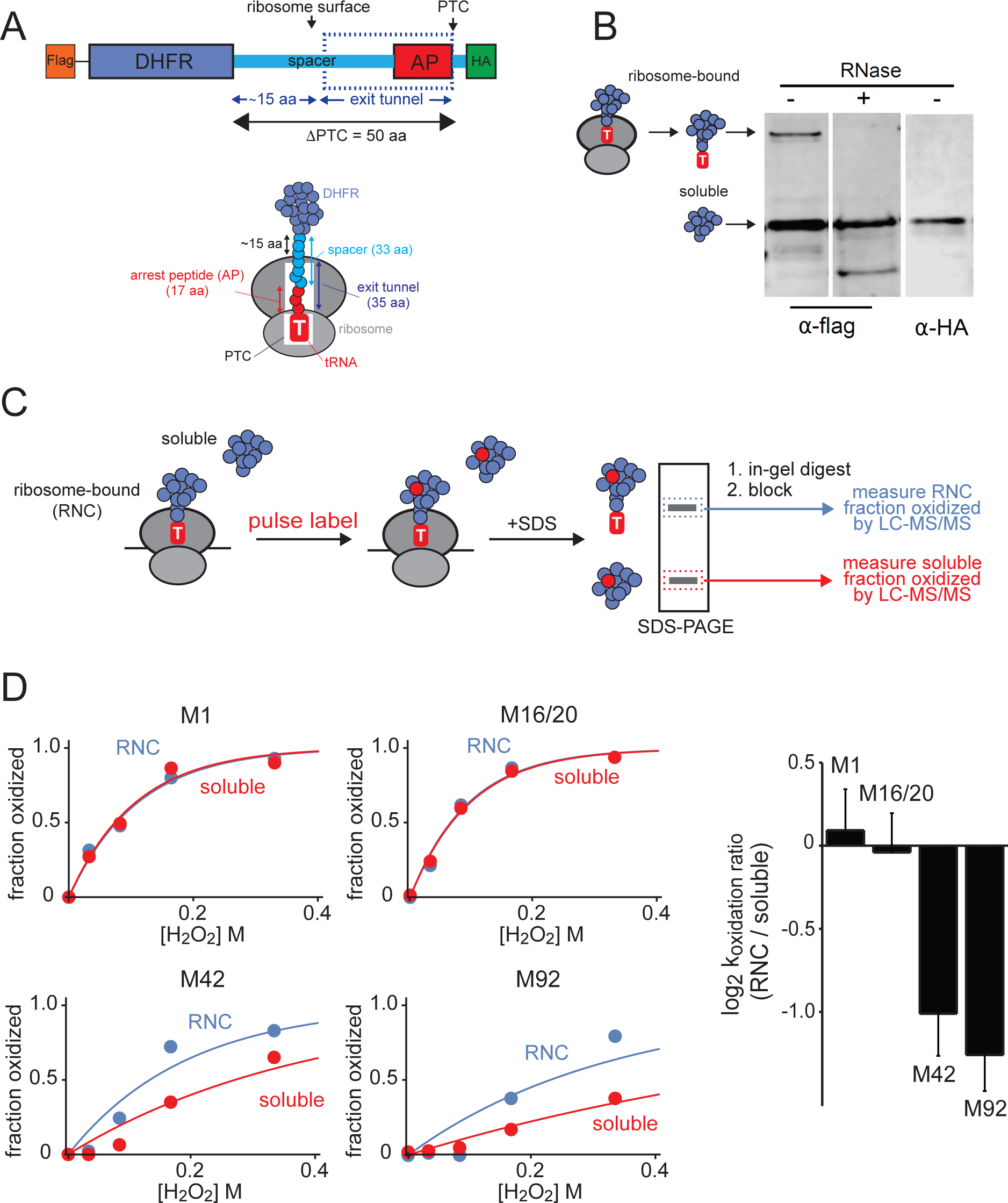
Methionine oxidation kinetics within DHFR-AP (ΔPTC = 50aa) in ribosome-bound (RNC) and soluble forms. **A)**In DHFR-AP constructs, DHFR is fused with the arrest peptide (AP) from SecM separated by an unstructured spacer. In the initial experiments shown in this figure, the distance between the C-terminus of DHFR and the ΔPTC is expected to be 50 aa, creating a distance of 15 aa between DHFR and the ribosome surface. The construct was epitope-tagged FLAG and HA at the N- and C-termini, respectively. The dashed box indicates the expected position of the ribosomal exit tunnel relative the arrested construct. **B)** As expected, Western blots indicate that tRNA-bound DHFR-AP generated by AP-mediated translational arrest can be detected by anti-FLAG and not anti-HA and undergoes a size shift upon treatment with RNase A. **C)** Workflow for the mass spectrometric analysis of oxidized DHFR-AP. Upon *in vitro* translation of DHFR-AP, RNC and soluble forms of the protein accumulate and are subsequently oxidized with varying concentrations^18^O H_2_O_2_. The two forms of the protein are separated by SDS-PAGE by size. RNC (top) and soluble (bottom) bands were separately cut from the gel, digested with trypsin, blocked with ^16^O H_2_O_2_ and analyzed by LC-MS/MS to quantify fractional oxidation levels for methionine-containing peptides. **D)** Fractional oxidation and relative k_oxidation_ measurements of DHFR-AP (ΔPTC = 50-aa) methionines in RNC (blue) and soluble (red) forms. Curve fitting and error measurements are as described in Figure 1. See also Figure S2.

We expressed DHFR-AP in an *E. coli in vitro* translation system and probed the resulting mixture with anti-FLAG and HA antibodies (Figure 2B). The immunoblots indicated the presence of fully synthesized soluble DHFR-AP with the expected molecular weight (MW), as well as a high MW band which we hypothesized to be tRNA-bound DHFR generated by its translational arrest. The accumulation of peptidyl-tRNAs by AP-mediated translational arrest had been observed previously^18, 45^. We verified that the high MW band is indeed DHFR-tRNA by demonstrating its sensitivity to RNase A digestion and lack of detection of its C-terminus HA tag. Thus, expression of DHFR-AP in an *in vitro* translation system results in the steady-state accumulation of soluble DHFR that has completed synthesis as well as translationally arrested tRNA-bound DHFR - two species that can be readily separated by gel electrophoresis.

To compare the stabilities of soluble and ribosome-bound forms of DHFR-AP, we translated the fusion constructs *in vitro* and then exposed the resulting mixture to varying concentrations of ^18^O-labeled H_2_O_2_, blocked the remaining unoxidized methionines with ^16^O-labeled H_2_O_2_ after denaturation, separated fully synthesized and translationally arrested DHFR-AP by SDS-PAGE and quantified fractional oxidation of each form by LC-MS/MS after in-gel trypsin digest (Figure 2C). In this way, we were able to compare the oxidation kinetics and folding stabilities of soluble and ribosome-bound DHFR-AP in parallel measurements within the same experimental sample. The data indicated that buried methionines (but not solvent-exposed methionines) had a faster rate of oxidation within ribosome-bound DHFR-AP (Figure 2D). Thus, ribosome-bound DHFR-AP appears to be less thermodynamically stable than its soluble form.

### Distance and salt dependence of ribosome-induced destabilization

We next quantified ribosome-induced destabilization in DHFR-AP constructs where DHFR had varying proximities to the ribosome. The constructs were created with variable spacer lengths such that the distance between the C-terminus of DHFR and the ΔPTC had a range of 34 to 70 amino acids. If the spacer has a fully extended conformation within the exit tunnel, DHFR would be expected to have scarcely cleared the exit tunnel in the 34 aa ΔPTC construct, and be separated from the ribosomal surface by 36 amino acids in the 70 aa ΔPTC construct. Oxidation experiments indicated that the elevated oxidation rates of buried methionines (M42 and M92) within ribosome-bound DHFR diminishes as a function of distance from the ribosome and is undetectable for the 70 aa ΔPTC construct (Figure 3). This trend was not detected for the exposed methionines (Figure S3). The results suggest that the destabilizing effect of the ribosome on nascent chains is strongly distance-dependent.

**Figure 3.**
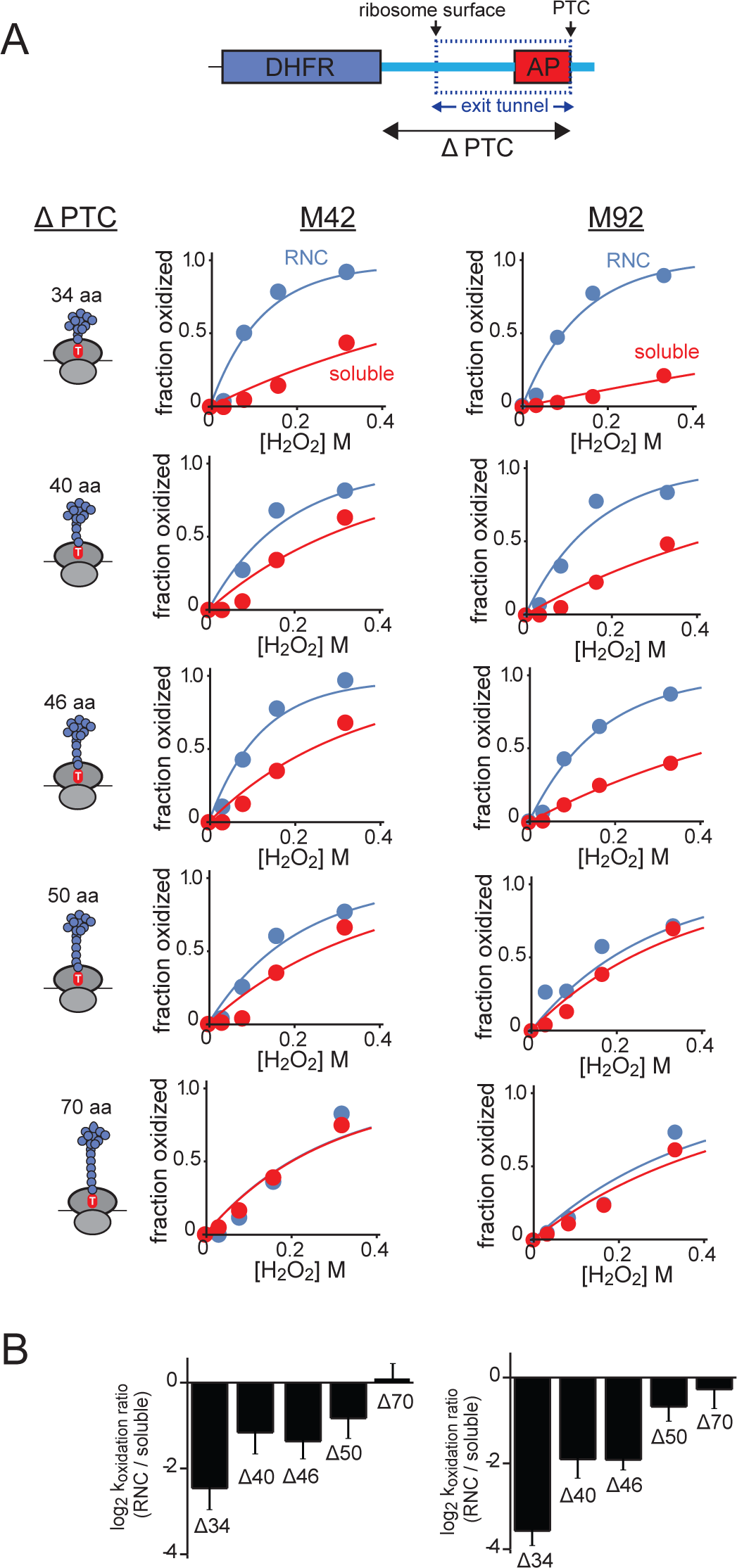
Distance dependence of ribosome-induced stability effects. **A)**Generation of DHFR-AP constructs with varying ΔPTC distances. Note that FLAG and HA epitope tags were removed from these and future constructs. B) Fractional oxidation and relative k_oxidation_ measurements of DHFR-AP buried methionines (M42 and M92) for constructs with variable ΔPTC lengths. Data corresponding to RNC and soluble forms of the proteins are indicated by red and blue colors, respectively. Curve fitting and error measurements are as described in Figure 1. See also Figures S2 and S3.

To determine whether electrostatic interactions play a role in the destabilizing effects of the ribosome on nascent chains, we repeated our experiments in the presence of increasing concentration of potassium chloride (KCl). Differences in oxidation levels of M42 and M92 observed between soluble and ribosome-bound DHFR at a constant H_2_O_2_ concentration (160 mM) diminished as a function of KCl concentration and were undetectable at 0.8 M KCl (Figure 4). The results suggest that electrostatic interactions contribute significantly to the ribosome’s ability to destabilize nascent DHFR.

**Figure 4.**
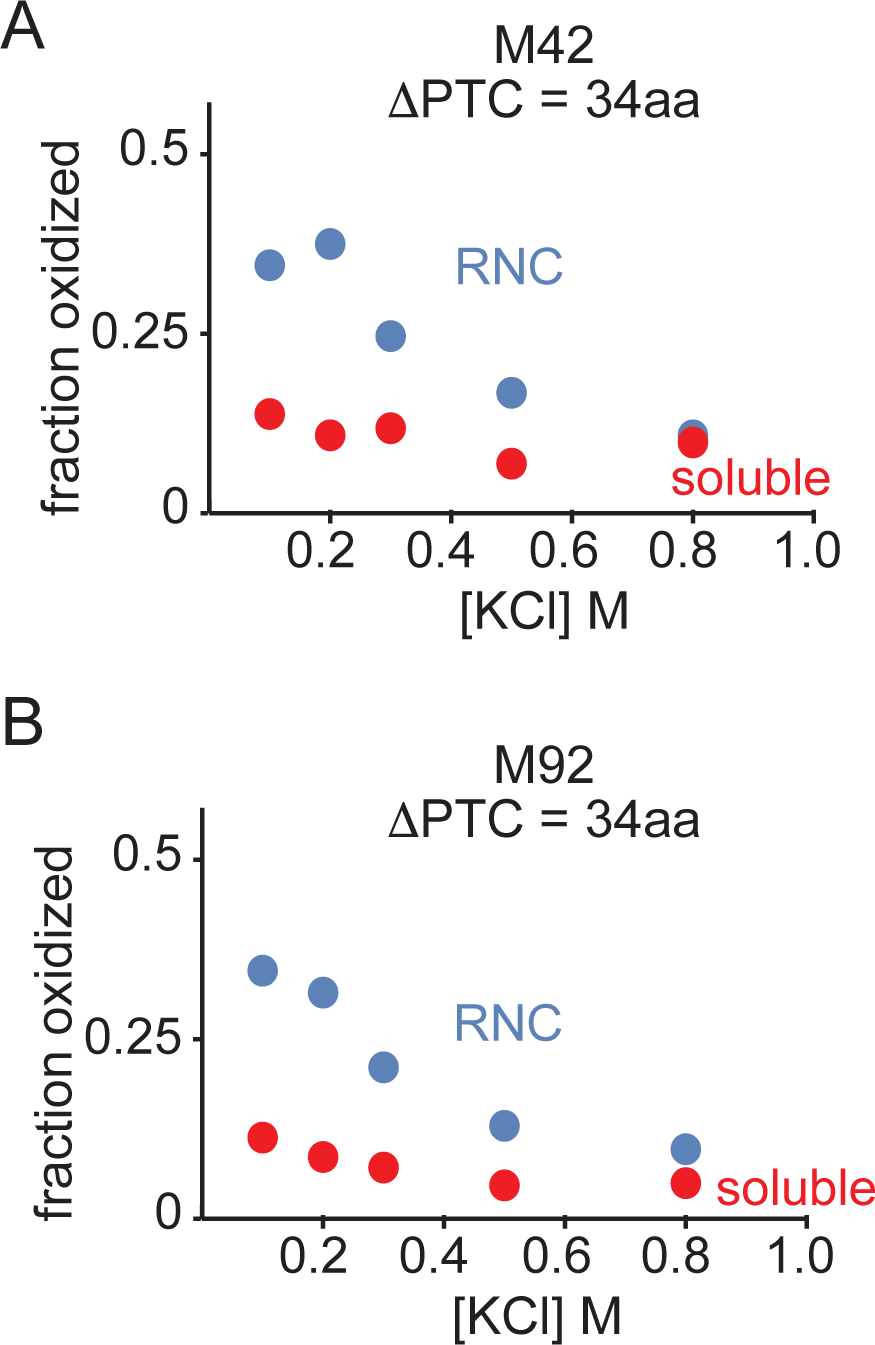
Salt dependence of ribosome-induced stability effects. Fractional oxidation measurements of buried methionines (M42 and M92) within DHFR-AP (ΔPTC = 34 aa) as function of KCl concentrations. Proteins were all labeled with 0.16 M H_2_O_2_. Data corresponding to RNC and soluble forms of the proteins are indicated by red and blue colors, respectively.

### Ligand binding by ribosome-bound DHFR

The above data indicate that the ribosome destabilizes nascent DHFR, hence diminishing the free energy difference between conformations that protect methionines from oxidation and conformations that expose them to oxidation. However, it remained unclear whether the protected state of ribosome-bound DHFR is conformationally equivalent to the native state of soluble DHFR. In other words, the observed ribosome-induced destabilization of DHFR maybe be a result of altered energetics of its native structure, or it may be due to the fact that the structure of ribosome-bound DHFR is distinct from its soluble form in a way that exposes methionines to oxidation. To distinguish between these two possibilities, we compared the oxidation kinetics of DHFR-AP in the presence and absence of MTX (Figure 5). The data indicate that MTX reduces the oxidation rates of buried methionines within both soluble and ribosome-bound DHFR-AP. Thus, ribosome-bound DHFR has a native-like conformation that is capable of binding MTX and undergoing ligand-induced stabilization. This observation is consistent with the idea that ribosome-bound DHFR has a native-like conformation with decreased thermodynamic stability.

**Figure 5.**
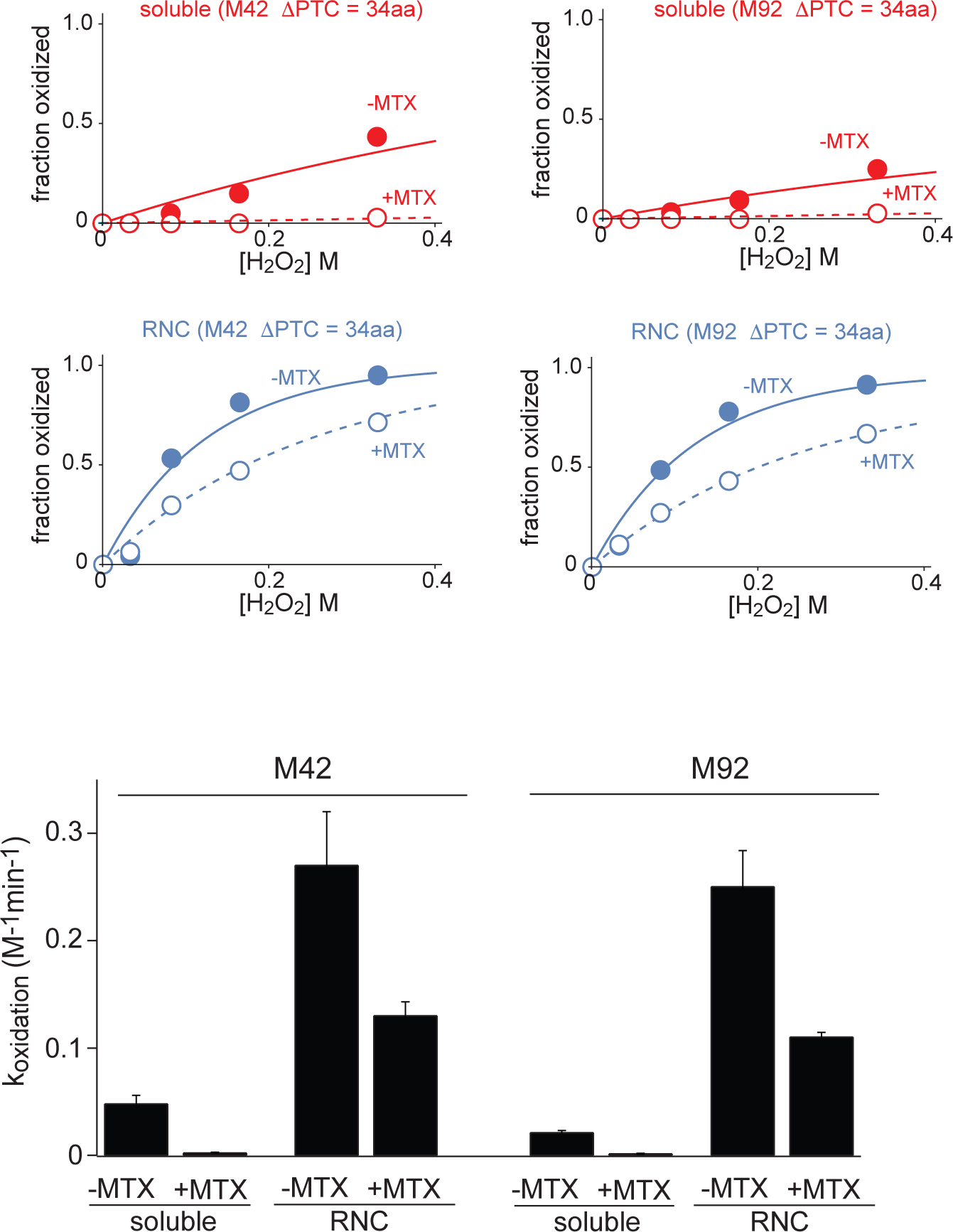
Ligand binding by ribosome-bound (RNC) and soluble DHFR-AP. Fractional oxidation and k_oxidation_ measurements of DHFR-AP buried methionines (M42 and M92) in the presence and absence of 300 µM MTX. Data corresponding to RNC and soluble forms of the proteins are indicated by red and blue colors, respectively. Curve fitting and error measurements are as described in Figure 1.

### Methionine oxidation kinetics in ribosome-bound CheY and DinB

To evaluate whether ribosome-induced effects observed for nascent DHFR extends to other proteins, we analyzed two additional model proteins: *E. coli* chemotaxis protein Y (CheY) and DNA polymerase IV (DinB) (Table 1). CheY is a single-domain protein similar in size to DHFR (129 aa). Like DHFR, it is an acidic protein with a relatively low isoelectric point (pI=4.9) and is thus negatively charged at physiological pH^48^. DinB is a larger (351 aa) basic protein with a higher isoelectric point (pI=8.5) and is positively charged at physiological pH. Both proteins have a number of buried and solvent-exposed methionines (Figure 6, S4). We generated AP fusions of the two proteins with spacer lengths that resulted in ΔPTC of 40 amino acids. Similar to DHFR, *in vitro* translation of the two constructs resulted in the accumulation of high MW tRNA-bound arrested nascent proteins that could be separated from the fully-synthesized forms of the proteins by SDS-PAGE, allowing us to compare the oxidation kinetics of the soluble and ribosome-bound forms of the proteins (Figure 6, S4).

**Figure 6.**
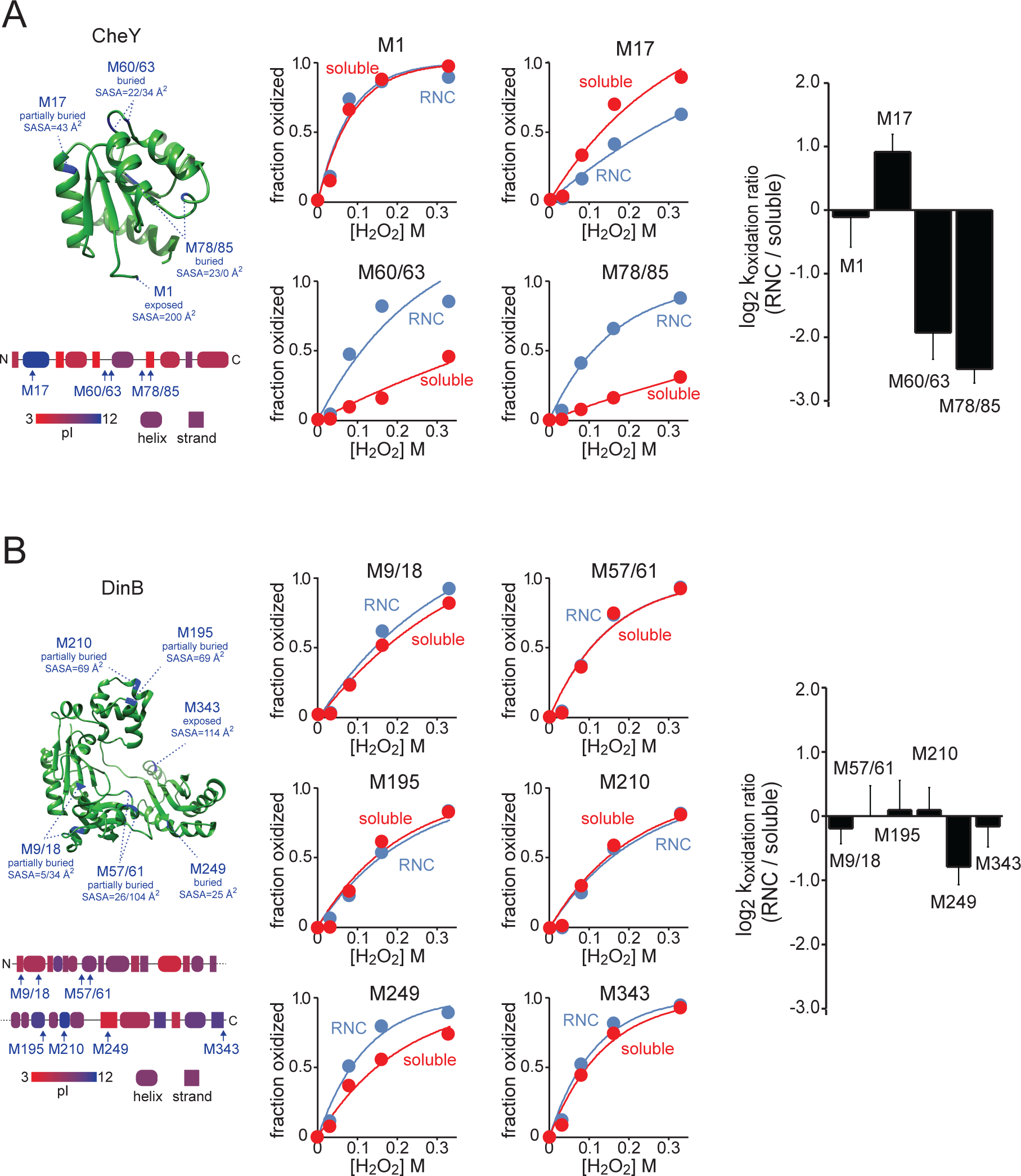
Methionine oxidation kinetics within CheY-AP and DinB-AP (ΔPTC = 40aa) in ribosome-bound (RNC) and soluble forms. Fractional oxidation and relative k_oxidation_ measurements of CheY-AP (P0AE67) and DinB-AP (Q47155). The constructs have ΔPTC of 40 aa. For the two proteins, structures, locations of methionines, SASA measurements and pI distributions are illustrated as in Figure 1. CheY M60/M63, M78/85 and DinB M9/18, M57/61 are nearby pairs of methionines that are contained in the same tryptic peptide. Data corresponding to RNC and soluble forms of the proteins are indicated by red and blue colors, respectively. Curve fitting and error measurements are as described in Figure 1. See also Figure S4.

**Table 1.** Measured differences in folding stabilities between RNC and soluble forms of DHFR, CheY and DinB.

| protein | gene | uniprot | length<br>(aa) | pI | charge | methionines | ΔPTC<br>(aa) | $\Delta\Delta G_{\text{RNC-soluble}}$<br>(kcal/mol) |
| --- | --- | --- | --- | --- | --- | --- | --- | --- |
| dihydrofolate reductase<br>(DHFR) | folA | P0ABQ4 | 159 | 4.8 | -10.1 | M42 | 34 | $1.05 \pm 0.31$ |
| | | | | | | | 40 | $0.45 \pm 0.31$ |
| | | | | | | | 46 | $0.62 \pm 0.25$ |
| | | | | | | | 50 | $0.35 \pm 0.29$ |
| | | | | | | | 70 | $0.00 \pm 0.27$ |
| | | | | | | M92 | 34 | $1.53 \pm 0.21$ |
| | | | | | | | 40 | $0.81 \pm 0.27$ |
| | | | | | | | 46 | $0.82 \pm 0.15$ |
| | | | | | | | 50 | $0.29 \pm 0.21$ |
| | | | | | | | 70 | $0.12 \pm 0.28$ |
| chemotaxis protein Y<br>(CheY) | cheY | P0AE67 | 129 | 4.9 | -4.0 | M17 | 40 | $-0.39 \pm 0.17$ |
| | | | | | | M60/63 | 40 | $0.83 \pm 0.25$ |
| | | | | | | M78/85 | 40 | $1.07 \pm 0.13$ |
| DNA polymerase IV<br>(DinB) | dinB | Q47155 | 351 | 9.3 | 8.5 | M9/18 | 40 | $0.09 \pm 0.14$ |
| | | | | | | M249 | 40 | $0.34 \pm 0.17$ |

Within CheY-AP, the buried or partially buried methionines M60, M63, M78 and M85 oxidize significantly faster on the ribosome compared to their soluble forms (Figure 6A). This suggests that, much like DHFR, the regions of CheY that harbor these methionines are destabilized by the ribosome. In contrast, the oxidation of M17 is significantly slower in the ribosome-bound form of CheY. It is important to note that although CheY is an acidic protein with an overall net negative charge, M17 resides in a secondary structure element (helix 1) that has pI of 11.8 and a locally high net positive charge. The results indicate that the ribosome does not have a uniformly destabilizing effect on all folding domains, and may in fact have the ability to locally stabilize positively charged regions of nascent proteins.

Within DinB-AP, the buried or partially buried methionines M9, M18, M57, M61, M195 and M210 have very similar oxidation rates on and off the ribosome suggesting that the stability of regions harboring these methionines is unaffected by the ribosome (Figure 6B). These regions are distributed in basic regions of the protein that have a marginally high pI. In contrast, the buried M249 residue resides in an acidic beta strand secondary structure element has a local negative charge. This residue has a faster oxidation rate on the ribosome indicative of localized ribosome-induced destabilization.

### Magnitude of stability changes (ΔΔ_Gfolding_) induced by the ribosome

Above analyses of DHFR, CheY and DinB indicate that the effect of the ribosome on the folding stability of nascent polypeptides is variable between proteins, and within different regions of the same protein. Specifically, in these three proteins, the destabilizing effects of the ribosome are most pronounced for acidic regions whereas basic regions are unaffected or stabilized. By making some simplifying assumptions regarding the nature of the methionine oxidation reaction, the differences in oxidation rates between the ribosome-bound and soluble forms of proteins can be used to approximate the difference in the free folding energy between the two. The details of the kinetic model used for this analysis has been described previously^3, 37^ and outlined in the Methods section below. Briefly, if we make the assumptions that 1) rate of folding is greater than the rate of unfolding (i.e. the protein is predominantly populating the folded state relative to the unfolded state), 2) rate of folding is greater than the rate of oxidation (i.e. the folding reaction is a rapid pre-equilibrium relative to the oxidation reaction) and 3) the oxidation reaction is occurring predominantly from the unfolded state (i.e. methionine is largely protected from oxidation in the folded state), then:

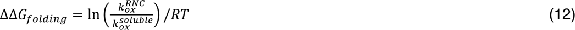

Where 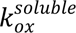 and 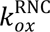 are the observed rate constants of oxidation in the soluble and ribosome-bound forms of the protein and ΔΔ_Gfolding_ is the difference in the folding stabilities (ΔΔ_Gfolding_) of the two forms. In this context, ΔΔ_Gfolding_ refers to the stability of the localized folding unit in the protein whose unfolding is the predominant route of solvent exposure for the residue (See also *Protein folding model and measurement of ΔΔ_Gfolding_ in* Methods). Measured ΔΔ_Gfolding_ values for the methionines analyzed in this study are plotted in Figure 7 and listed in Table 1.

**Figure 7.**
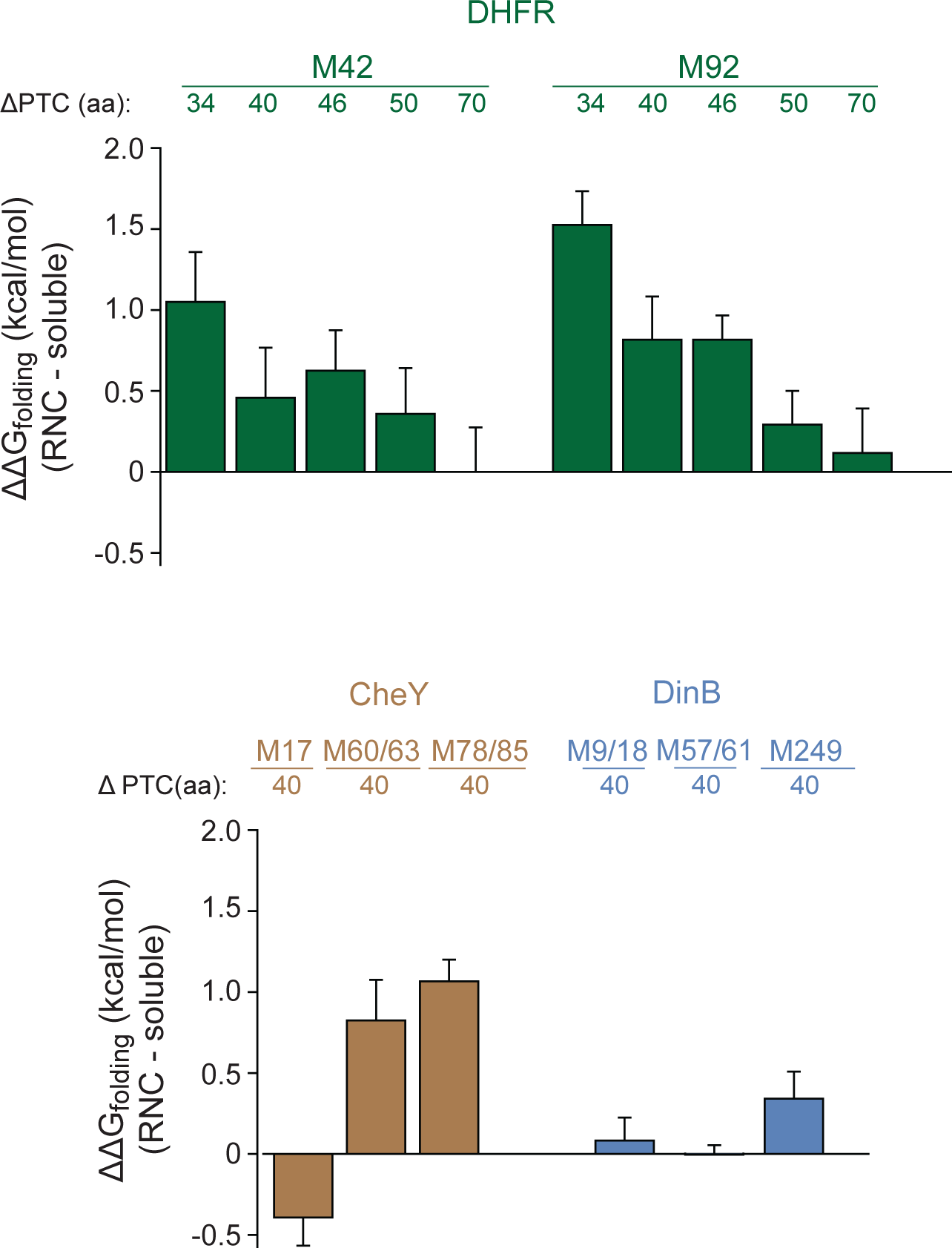
Thermodynamic folding stability changes induced by the ribosome (ΔΔ_Gfolding_) within the analyzed model protein. Differences in oxidation kinetics of buried and partially buried (SASA < 50 Å^2^) methionines between RNC and soluble forms of the analyzed proteins were converted to ΔΔG measurements in accordance to the two-state model outlined in the manuscript.

## Discussion

Previously, the dynamics and folding of RNCs had been studied by a number of methodologies including optical tweezers, NMR spectroscopy and limited proteolysis ^8, 16, 20, 21^. Each of these approaches offers unique methodological advantages and some limitations. NMR spectroscopy allows for analysis of residue-specific dynamics, but is typically not practical for analysis of diverse polypeptides, particularly larger proteins. Optical tweezers are challenging to implement and lack the ability to provide localized folding stability information. Pulse proteolysis under denaturing conditions can measure the global folding stabilities of ribosome-bound model proteins. However, this approach requires the addition of molar concentrations of chemical denaturants to the RNC complex which can influence ribosome-induced thermodynamic effects. The requirement for denaturation also limits the analysis to unstable proteins that require low concentrations of denaturants for their unfolding.

The bottom-up proteomic approach described in this study combines some of the benefits of the above approaches and offers a number of novel advantages. First, the method allows for quantitative measurements of thermodynamic folding stabilities under native conditions without denaturant-induced unfolding of nascent peptides. Second, the approach can be used to measure sub-global folding stabilities within localized regions of proteins. Third, the analysis can be applied to a range of proteins regardless of size and folding stability and allows for measurement of RNC folding stabilities within complex mixtures and in the presence of excess levels of other proteins and factors. However, the application of SPROX to RNCs has at least two significant limitations. First, the analysis is limited to protein regions that contain buried methionine residues. Second, methionine oxidation is not a benign modification and may in itself impact folding stability. Thus, oxidation of one methionine residue could artifactually alter the localized stability of a second methionine located elsewhere in the same protein. Although this possibility can not be discounted for any one protein, previous proteomic experiments have indicated that such scenarios are relatively rare and methionine oxidation rates provide relatively accurate measures of folding stability^49^.

Our results demonstrate that ribosomes can strongly modulate the folding stability of nascent polypeptides. The magnitude and direction of this effect appear variable between proteins - and even within different regions of the same protein. Our analyses of DHFR, CheY and DinB indicate that the ribosome has a tendency to destabilize acidic, negatively charged regions of nascent polypeptides, while basic regions are either unimpacted or stabilized (as in the case of helix 1 of CheY). The magnitude of ribosome-induced stability modulations were shown to be distance and salt dependent, thus implicating electrostatic interactions between the surface of the ribosome and nascent polypeptides as the possible cause of the observed stability effects. Because of the presence of ribosomal RNAs, the surface of the ribosome is a highly negatively charged environment. During the course of translation, electrostatic interactions between the ribosome surface and nascent polypeptides may provide a mechanism for altering the free energy difference between folded and unfolded conformations of localized folding domains. These results support previous data indicating that specific interactions between the ribosome surface and nascent chains can influence co-translational folding^16^.

In theory, the ribosome can alter the free energy of a protein’s folded state relative to its unfolded state either by disrupting the native structure of the protein, or by changing its energy relative to the unfolded state (while retaining the native conformation). In the case DHFR, we showed that the ribosome-bound protein has the ability to bind methotrexate. This observation suggests that ribosome-bound DHFR retains its ability to fold into its soluble native conformation. Thus, ribosomes can influence the energetics of native states without fundamentally altering their structures.

By making some simplifying approximations and employing a two-state protein folding model, we quantitatively measured the differences in thermodynamic folding stabilities of soluble and ribosome-bound proteins. These differences ranged from -0.4 to +1.5 kcal/mol depending on the model protein and its distance from the ribosome surface. The ΔΔG values measured for DHFR were +1.1 and +1.5 kcal/mol for M42 and M92, respectively. These measurements are slightly lower than ∼+2 kcal/mol stability differences measured for DHFR by pulse proteolysis^21^. However, it is important to note that whereas pulse proteolysis reports on the global stabilities of proteins, the SPROX methodology provides measurements of local/sub-global stabilities (see also Experimental Procedures). Similar to what has been observed in native-state hydrogen-deuterium exchange (HDX) experiments^50^, solvent exposure of most protein regions can occur by localized unfolding events that are more energetically favorable than global unfolding. Thus, localized thermodynamic folding stabilities of most protein regions are expected to be lower and differentially impacted by the ribosome in comparison to global folding stabilities.

What could be the functional significance of ribosome-induced stability effects on nascent polypeptides? As has been pointed by a number of previous studies^8^, the ribosome may alter the energetic landscape of nascent proteins in order to facilitate co-translational folding. For example, by destabilizing incompletely synthesized proteins, the ribosome may prevent premature folding and formation of detrimental folding intermediates until the entirety of a folding domain has emerged from the ribosomal exit tunnel. A second possibility is that by altering the folding stability of proteins, the ribosome can modulate rates of co-translational modification. Nascent proteins can be enzymatically modified during the course of their synthesis through processes that include deformylation, acetylation, hydrolysis of the initiator methionine and ubiquitination^51, 52^. By altering the stability of nascent peptides, the ribosome can enhance or restrict access to specific regions of proteins and influence the efficiency of enzymatic modifications. Determining the detailed relationships between ribosome-induced stability effects and co-translational folding and modifications will require further experimentation. The approach described in this study may facilitate such experiments by providing a path for conducting co-translational folding studies on proteome-wide scales.

## Methods

### Design and engineering of SecM arrested constructs

Expression plasmids contained genes for the three model proteins (folA, cheY and dinB) downstream of a T7 promoter and Shine-Dalgarno sequences. These genes were fused with the 17 amino acids arrest peptide from *E. coli* SecM with spacers of various lengths as described in the manuscript. The spacers were derived from unstructured domains of the protein LepB as described previously^53^. The exact sequences of the model protein-AP fusions are illustrated in Supp Figure S2. In the case of the initial validation experiments illustrated in Figure 2, FLAG and HA epitope tags were fused at the N- and C-termini of the DHFR-AP construct.

### In vitro translation, SDS-PAGE and Western blots

Coupled *in vitro* transcription and translation reactions were carried out using the *E. coli* PURExpress kit (New England Biolabs, Cat no. E6800). The components and buffer composition of this system have been described by Shimizu et al.^42^. Standard reactions had a volume of 25 µl and contained 1.5 µg DNA templates incubated at 37 ℃ for 30 minutes. In some experiments, synthesized proteins were labeled with 10 µCi ^35^S Methionine (Perkin Elmer). Following translation, 5 µl of the reactions were denatured with LDS sample buffer (Invitrogen) and heated at 65 ℃ for 10 minutes. The samples were subsequently run on 12% NuPAGE Bis-Tris Gel (Invitrogen) and transferred on PVDF membrane (BioRad). The membrane was then exposed on BioBlot film (LPS) overnight in dark at room temperature.

For Western blot experiments, anti-FLAG antibody (DYKDDDDK Tag Monoclonal Antibody, Dylight 680. Thermo, MA1-91878-D680) and anti-HA antibody (HA Tag Monoclonal Antibody, Dylight 800 4X PEG. Thermo, 26183-D800) were used. The primary antibodies were diluted 1:1000 ratio in intercept blocking buffer (Li-Cor) and incubated with membranes overnight at 4 ℃. The blot was imaged by Li-Cor imager (Odyssey). In the puromycin labeling experiment (Figure S2A), biotin-dC-puromycin (Jena Bioscience) was mixed with *in vitro* translation reaction to a final 10 µM. *E. coli* SecM was then translated at 37 ℃ for 1 hour. 5 µl of protein was denatured and used for Western Blot analysis. Streptavidin Antibody, Dylight 800 (Thermo Fisher) was used for fluorescence detection of puromycin tagged proteins.

### DHFR and DHFR-AP in vitro translation in the presence of MTX, GdmCl and KCl

For experiments carried on non-arrested WT DHFR (Figure 1), translation reactions were carried out for 4 hours at 37 ℃. For MTX treated samples, methotrexate (Sigma) was dissolved in DMSO to make 5 mM stock solution. 1.5 µl was added to a 25 µl *in vitro* translation reactions with final concentration 500 μM and incubated 1 hour prior to SPROX. For GdmCl treatments, guanidine hydrochloride (Sigma) was dissolved in water to make a 12 M stock solution. 5 μl was added to a 25 μl *in vitro* translation reaction with final concentration 2 M and incubated 1 hour prior to SPROX.

In translation reactions with DHFR-AP in the presence of MTX (Figure 5), the final concentration of MTX was 300 μM. For translation reactions with DHFR-AP in the presence of KCl (Figure 4), 25 ul reactions were divided into 4 aliquots, each containing 5 µl. The protein was pre-incubated with KCl with final concentration of 0.1 M, 0.2 M, 0.3 M, 0.5 M and 0.8 M at 37 ℃ for 10 minutes prior to SPROX.

### SPROX and MS Sample preparation

In a typical SPROX analysis, 25 μl *in vitro* translation reactions were incubated at 37 ℃ for 30 minutes and divided into 5 μl aliquots. Each aliquot was incubated with ^18^O H_2_O_2_ (Sigma) with final concentration of 0 M, 0.032 M, 0.08 M, 0.16 M and 0.32 M for 30 minutes at 37 ℃, the total volume of each tube was brought up to 10 μl using a buffer containing 10mM magnesium acetate and 100mM potassium chloride, pH 7.4. The oxidation was quenched by 2-fold molar concentrations of sodium sulfite (Sigma) for 5 minutes. Note that for the KCl titration experiments, only 0.16 M concentration of ^18^O H_2_O_2_ were used. Subsequently, the samples were mixed with 4X LDS sample buffer and denatured at 95 ℃ for 10 minutes. The denatured proteins were then run on 12% Bis-Tris gel to desalt and to separate tRNA-bound and soluble proteins. The gels were subsequently stained with Bio-safe Coomassie (BioRad) for 1 hour at room temperature and washed in distilled water overnight at 4 ℃. Individual gel bands were cut and diced in ∼1mm^3^ pieces. In cases where the tRNA-bound and soluble proteins bands could not be directly visualized in Coomasie-stained gels (because of low amounts), regions of gels corresponding to band sizes detected in ^35^S-labeled gels were excised for analysis.

Gel pieces were soaked and washed in 100 mM ammonium bicarbonate (Fluka) for 10 minutes using sufficient volume to cover all the pieces. The gel pieces were then washed in 50 mM ammonium bicarbonate with 50% acetonitrile (ACN) (Thermo Fisher) for 10 minutes and dehydrate in pure ACN. Disulfide bonds were reduced by adding 10 mM Dithiothreitol (DTT) (Fisher) and incubating at 55 ℃ for 1 hour. Protein alkylation reactions were then carried out with 50 mM Iodoacetamide (IAA) (Sigma) at room temperature for 30 minutes in the dark. The gel pieces were then washed and dehydrated with ammonium bicarbonate and ACN. Trypsin (Pierce) was added at a 1:100 ratio (trypsin/protein), incubating overnight at 37℃. Trypsin was quenched by adding 0.1% trifluoroacetic acid (TFA) (Pierce) with 50% ACN. Supernatant was collected and lyophilized. To block methionine residues, final concentration of 0.16 M of ^16^O H_2_O_2_ solution was added to resuspend peptide and incubate at 37 ℃ for 30 minutes. To remove excess H_2_O_2_, the peptide was loaded on homemade C18 spin columns. Peptides were washed with 0.1% TFA twice and eluted with 0.1% TFA in 50% ACN, lyophilized and resuspended in 25 µl of 0.1% TFA.

### Mass spectrometric analysis

Peptides were injected onto a homemade 30 cm C18 column with 1.8 µm beads (Sepax), with an Easy nLC-1200 HPLC (Thermo Fisher), connected to a Fusion Lumos Tribrid mass spectrometer (Thermo Fisher). Solvent A was 0.1% formic acid in water and solvent B was 0.1% formic acid in 80% acetonitrile. Ions were introduced to the mass spectrometer using a Nanospray Flex source operating at 2 kV. The gradient was started at 3% B and held for 2 minutes, increased to 10% B over 5 minutes, increased to 38% B over 38 minutes, then ramped up to 90% B in 3 minutes and was held for 3 minutes, before returning to starting conditions in 2 minutes and re-equilibrating for 7 minutes for a total run time of 60 minutes. The Fusion Lumos was operated in data-dependent mode, with MS1 scans acquired in the Orbitrap, and MS2 scans acquired in the ion trap. The cycle time was set to 1.5 seconds. Monoisotopic Precursor Selection (MIPS) was set to Peptide. The full scan was done over a range of 375-1400 m/z, with a resolution of 120,000 at m/z of 200, an AGC target of 4e5, and a maximum injection time of 50 ms. Peptides with a charge state between 2-5 were picked for fragmentation. Precursor ions were fragmented by collision-induced dissociation (CID) using a collision energy of 30% with an isolation width of 1.1 m/z. The Ion Trap Scan Rate was set to Rapid, with a maximum injection time of 200 ms, an AGC target of 1e4. Dynamic exclusion was set to 20 seconds.

### Database search

Raw files for all samples were searched against the *E. coli* K12 UniProt database (UP000000625_83333, downloaded 02/16/2022) using MSFragger and Fragpipe software^54^. Peptide and protein quantifications were performed with Fragpipe default parameter settings. ^18^O, ^16^O, and N-terminal formylation were set as variable modifications; carbamidomethyl cysteine was set as a fixed modification. MS1 spectra from raw files were converted to .ms1 format using MSConvert software^55^. The PSM search results from Fragpipe and the MS1 spectra were used to determine the fraction oxidized for each methionine-containing peptide.

### Measurement of surface accessible surface area (SASA)

SASA measurements were computed for *E. coli* available proteins as previously described^37^. Briefly, solvent accessible surface areas of methionine residues were calculated in Pymol, with dot density set to 3. SASA values for *E. coli* were computed using structures available from the AlphaFold Protein Structure Database using AlphaFold2^56^. The top five ranked models for each protein were then used to calculate SASAs as above, keeping only models with an average plDDT value >70.

### Measurement of fractional oxidation and oxidation rate constants

MS1 spectra matched to methionine-containing peptides were analyzed using a custom algorithm to quantify the ratio of intensities between ^16^O- an ^18^O-labeled peptides^39^ with slight modifications to account for changes in our experimental design (e.g. the use of ^18^O as pulse and ^16^O as block rather than *vice versa*) and the ability to analyze peptides with more than one methionine. In cases where multiple methionine residues were mapped to a single peptide, the spectra were fit to a model that assumes all methionines in the peptide have the same level of oxidation as described previously^57^. Thus, in these peptides, the reported oxidation level is the consensus value for all methionines that best fits the observed spectrum. In cases where multiple PSMs and charge states were identified for a given peptide, the highest intensity spectrum was quantified. ^16^O to ^18^O ratios (R_ox_) were converted to fractional oxidation (F_ox_) measurements using the following equation:

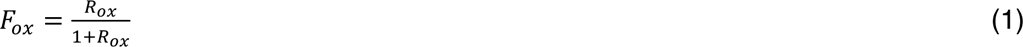

To measure oxidation rates for a given peptide, F_ox_ measurements were plotted as a function of [H_2_O_2_] and were fitted to the following equation:

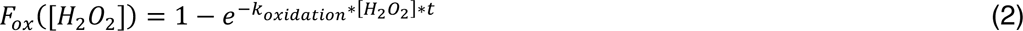

Where k_oxidation_ is the second order rate constant for methionine oxidation and t is the oxidation time (30 minutes). Here, *k_oxidation_*\*[H_2_O_2_] represents the pseudo first-order rate constant for the oxidation reaction as [H_2_O_2_] is in excess of protein-bound methionine concentrations. *k_oxidation_* values were determined using non-linear curve fitting based on the Levenberg-Marquardt algorithm. For plots displaying the fitted k_oxidation_ values, error bars represent the standard errors for parameter estimates based on the fitted model.

### Protein folding model and measurement of Δ_Gfolding_

To interpret methionine oxidation kinetics in terms of protein folding thermodynamics, we considered the following two-state folding model:

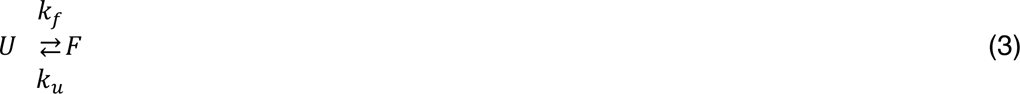

Where U and F are the folded (protected) and unfolded (unprotected) conformations of the protein, and *k_f_* and *k_u_* are the folding and unfolding rate constants, respectively, and thus:

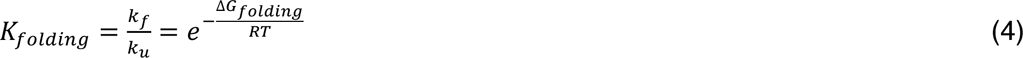

Where *K_folding_* is the folding equilibrium constant, *Δ_Gfolding_* is the thermodynamic folding stability (free energy difference between the folded and unfolded states), *R* is the gas constant, and *T* is the absolute temperature in Kelvin. Thus, the oxidation of methionines within proteins can be modeled as:

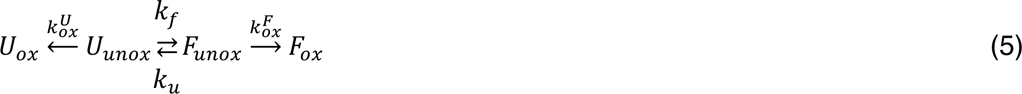

Where U_unox_ and F_unox_ are the folded and unfolded conformations of the protein harboring an unoxidized methionine residue, U_ox_ and F_ox_ are conformations harboring the oxidized methionine residue, and 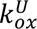 and 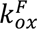 are the oxidation rate constants from the unfolded “4 ”4 and folded states, respectively. Akin to the kinetic model that is commonly used to interpret hydrogen-deuterium exchange (HDX) experiments^50^, the observed rate constant for oxidation can be expressed as:

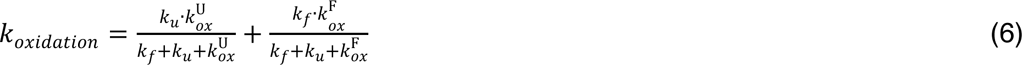

In a folded protein, *k_f_* >> *k_u_*, and hence:

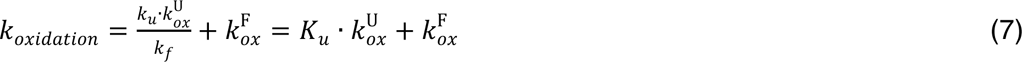

If we assume that a methionine is mostly protected from oxidation, then 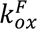 is negligible and:

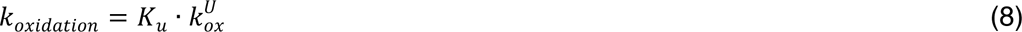

If we consider a protein that can exist in either RNC or soluble states, the ratio of oxidation rates in these two states is related to their respective folding equilibria as follows:

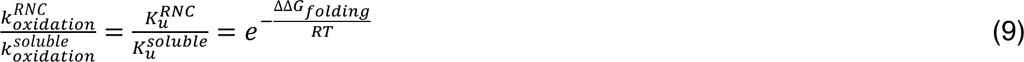

Where ΔΔ_Gfolding_ is the difference in folding free energies between the two states, R is the gas constant, and T is the absolute temperature in Kelvin.

For methionines that are readily oxidized from the folded state (i.e. exposed methionines), equation 5 simplifies to:

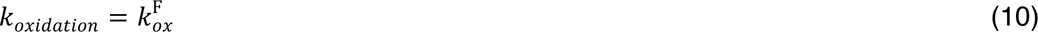

And hence:

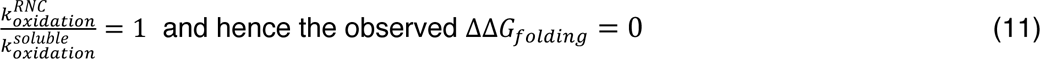

It is important to note that in the above model, “U” does not necessarily represent the globally unfolded protein but instead the lowest energy conformation that exposes the methionine to oxidation. Similar to models used to interpret HDX experiments, both global and sub-global unfolding events can potentially expose a protected residue to solvent^50^. Hence, the measured stabilities in the above analyses represent the Δ_Gfolding_ for the most energetically favorable unfolding equilibrium that results in the localized exposure of the methionine to solvent.

## Acknowledgement

All mass spectrometry experiments were conducted at the University of Rochester Mass Spectrometry Resource Laboratory. We thank Dr. Mark Dumont, Dr. Dmitri Ermolenko, Dr. Elizabeth Grayhack, Dr. Dragony Fu, Dr. Anne Meyer and members of the Ghaemmaghami Lab for helpful discussions and suggestions. We are also grateful to the Gorbunova lab for sharing some of the reagents used in this study. This study was supported by grants from the National Institute of Health (R35 GM119502, S10OD025242.)

## Data Availability

All raw and processed data are available in the included Supporting Information and at the ProteomeXchange Consortium via the PRIDE partner repository (accession number PXD037995). Currently, the data can be accessed with the username and password xaIpHm9s.

## Abbreviations

SASA: Solvent-accessible surface area
RNC: Ribosome nascent chains
AP: arrest peptide
PTC: peptidyl transferase center
LC-MS/MS: liquid chromatography-tandem mass spectrometry
HX: hydrogen/deuterium exchange
SPROX: Stability of Proteins from Rates of Oxidation
MObB: Methionine Oxidation by Blocking
DHFR: dihydrofolate reductase
GdmCl: guanidinium hydrochloride
MTX: methotrexate
MW: molecular weight
pI: Isoelectric point
DTT: dithiothreitol
IAA: iodoacetamide
TFA: trifluoroacetic acid
ACN: acetonitrile

## Author Information

Corresponding Author *. Phone 585-275-4829

## Author Contributions

The study concept was conceived by R.T. and S.G. Its detailed planning was performed with contribution from all authors. The project was performed by R.T., with assistance from M.H. The mass spectrometry was performed by K.W., K.S. and J.H. Data analysis was conducted by R.T. and S.G. The first draft of manuscript was written by R.T. and S.G.

## Notes

The authors declare no competing financial interest.

## Supplementary Figures

**Figure S1.**
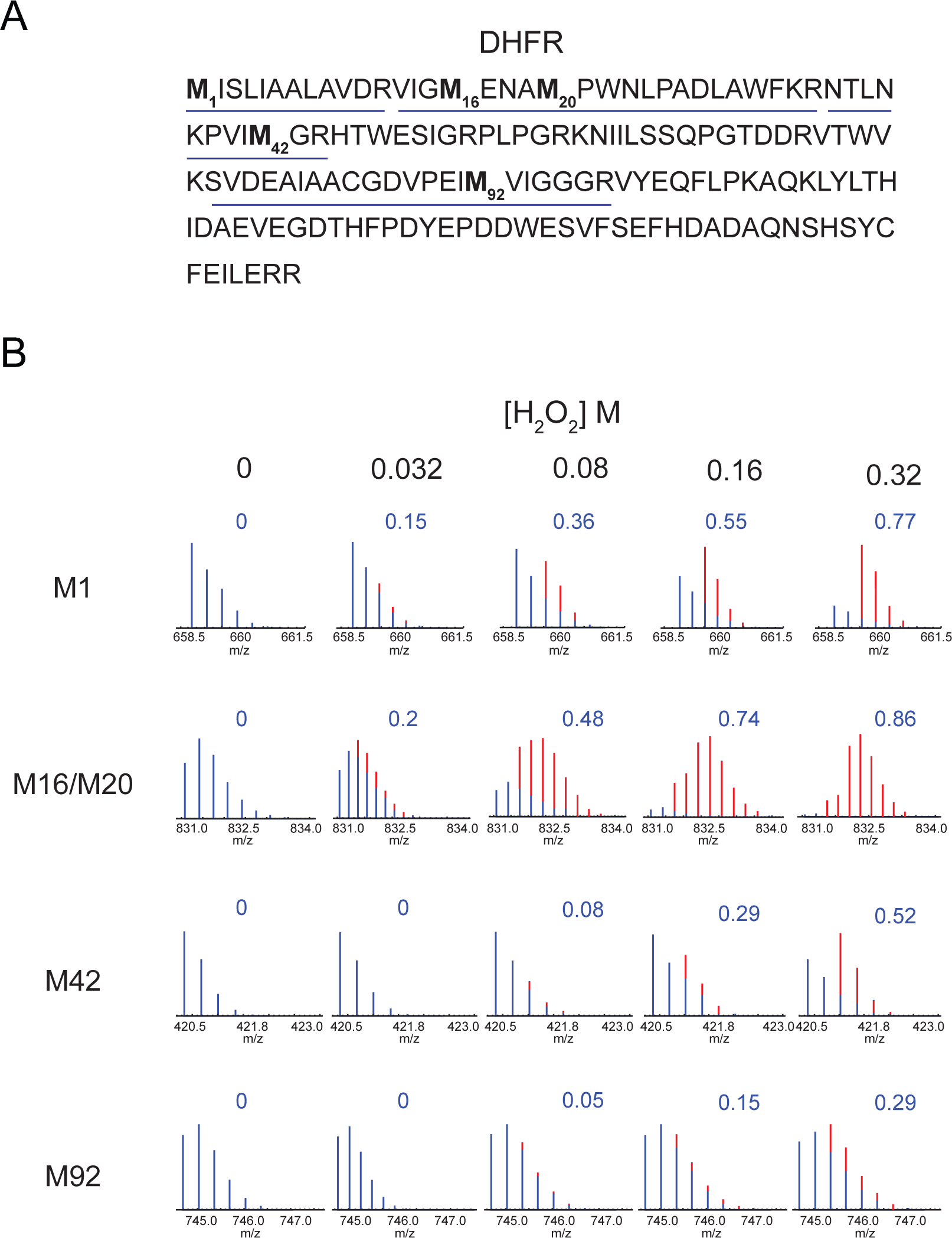
**A)** Protein sequence of *E.Coli* DHFR. Methionine-containing tryptic peptides that were detected by mass spectrometry and quantified are underlined. The analyzed methionines are highlighted in bold letters. **B)** Mass spectra of methionine-containing peptides detected by analysis of soluble full-length DHFR (See Figure 1C). Blue spectra indicate ^16^O oxidized peptides and red lines indicate ^18^O oxidized peptides. The calculated fractional oxidization levels determined from the intensity ratios of ^18^O/^16^O-oxidized spectra values are shown in blue. The spectra are shown as examples of raw data that were used to calculate methionine oxidation levels in experiments described in the remainder of the manuscript. Raw spectra for all experiments have been deposited in the PRIDE database (Acc: PXD037995)

**Figure S2.**
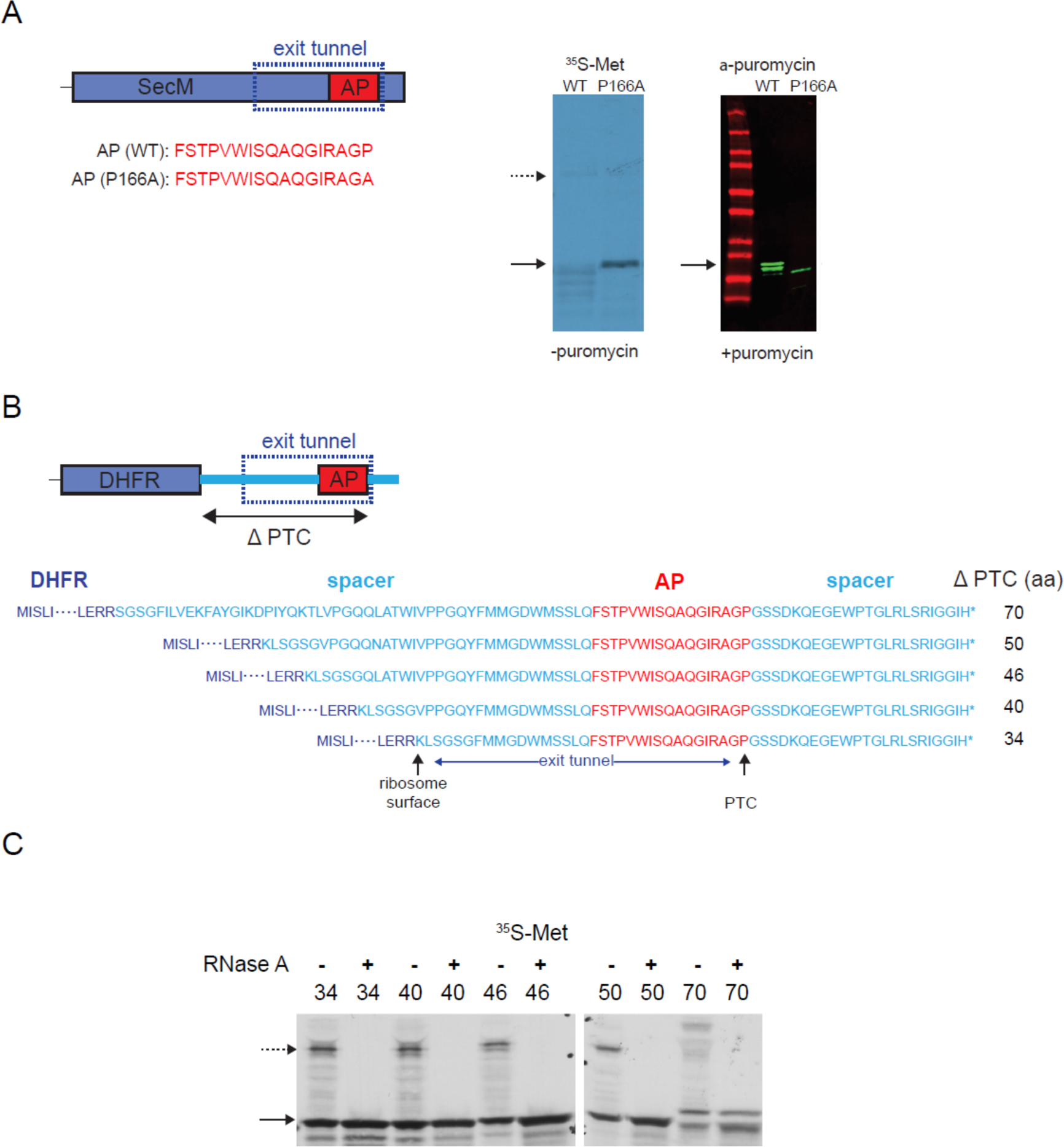
**A)**AP-mediated translational arrest of SecM. SecM protein contains a 17-aa arrest motif (AP) at its C-terminus that interacts with the ribosome exit tunnel and stalls translation. A P166A mutation in AP is known to relieve this pausing effect. AP-mediated translational arrest of SecM was verified in *in vitro* translation experiments. In ^35^S-methionine labeling experiment (left blot), the translation of SecM (bottom arrow) is inhibited in WT SecM and this arrest relieved in the P166A mutant. When translation of WT SecM is arrested, a higher MW band is observed (top arrow) that corresponds to the peptidyl-tRNA form of the protein that was observed in other AP fusion proteins (see for example Figure 2). The translational arrest of WT SecM can be relieved by addition of puromycin that releases the arrested SecM, which can then be detected by an anti-puromycin probe. **B)** Protein sequence of DHFR-AP constructs with ΔPTC values ranging from 34 to 70 aa. The arrest peptide sequence is highlighted in red and the spacer sequences are shown in light blue. The expected corresponding locations of the PTC, ribosome surface and the exit tunnel are shown by arrows. **C)** ^35^S-methionine labeled *in vitro* translation assays of DHFR-AP constructs with varying ΔPTC values indicating the formation of arrested RNase-sensitive peptidyl-tRNA forms of the proteins. The dashed arrow indicates the ribosome-associated forms and the solid arrow indicates soluble full length protein forms.

**Figure S3.**
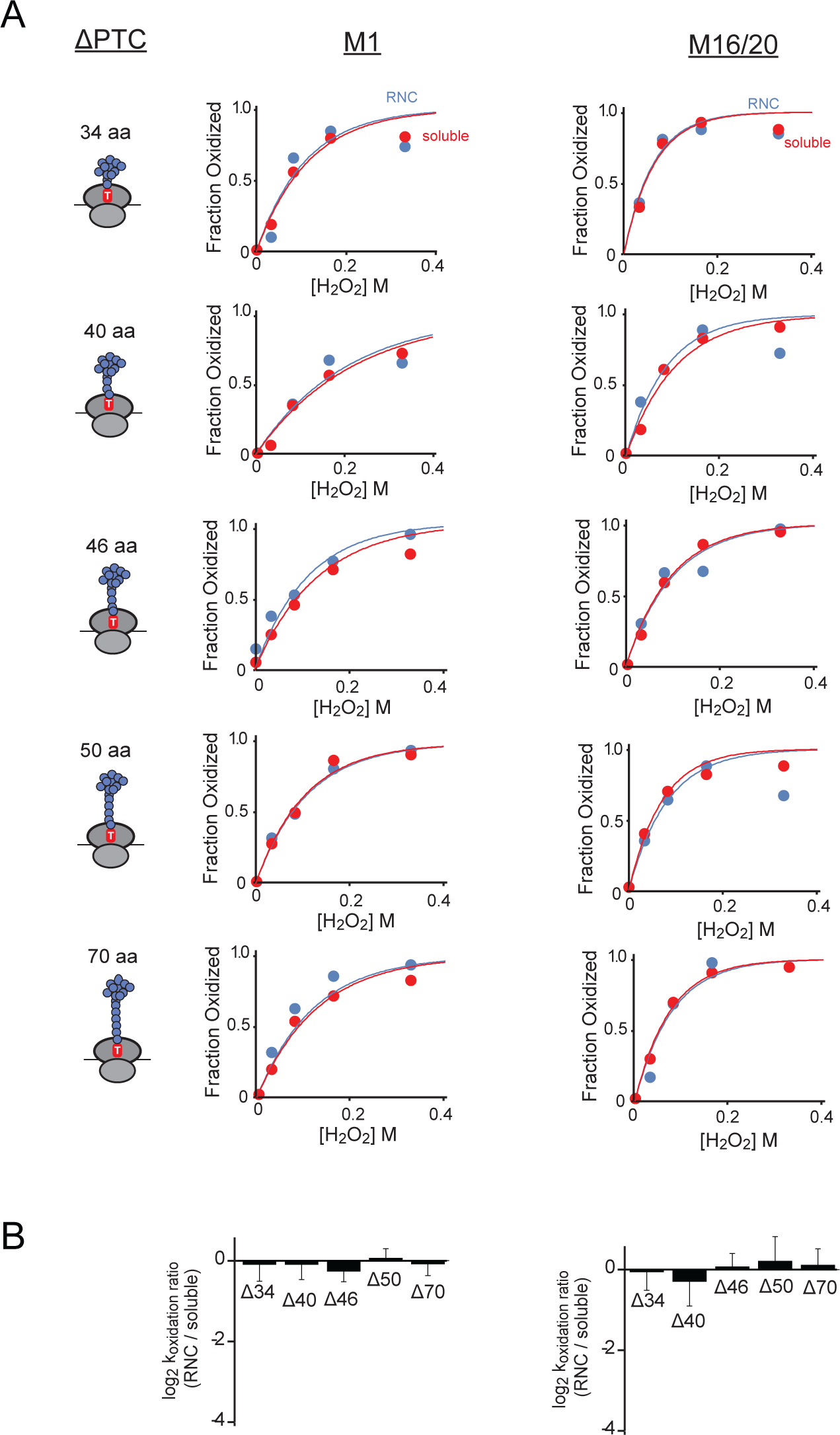
Fractional oxidation and relative k_oxidation_ measurements of DHFR-AP exposed methionines (M1 and M16/20) for constructs with variable ΔPTC lengths. Data corresponding to RNC and soluble forms of the proteins are indicated by red and blue colors, respectively. Curve fitting and error measurements are as described in Figure 1.

**Figure S4.**
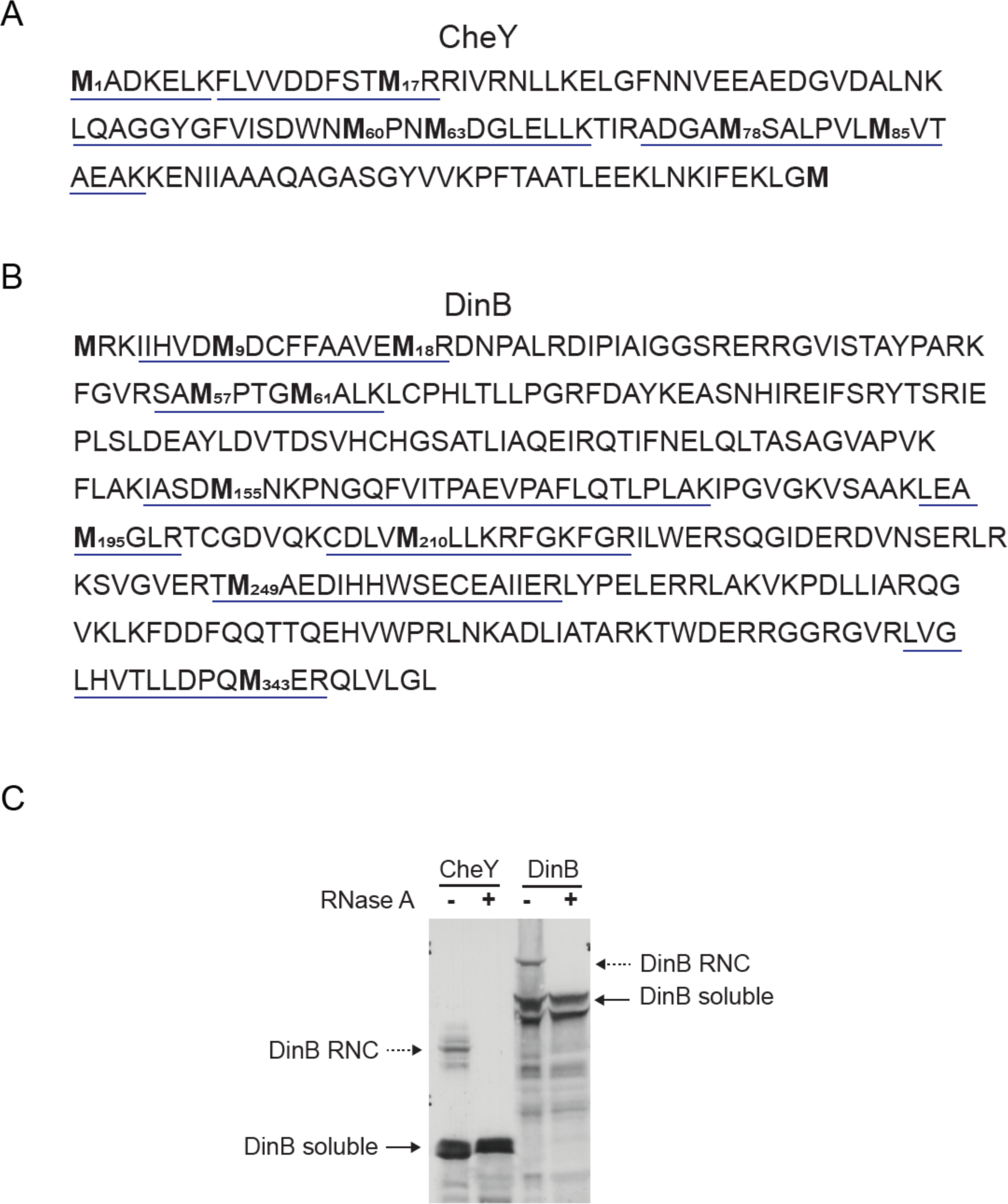
**A,B)** Protein sequence of *E.Coli* CheY (A) and DinB/Pol IV (B). Methionine-containing tryptic peptides that were detected by mass spectrometry and quantified are underlined. The analyzed methionines are highlighted in bold letters. **C)** ^35^S-methionine labeled *in vitro* translation assays of CheY-AP and DinB-AP constructs (ΔPTC = 40 aa) indicating the formation of arrested RNAse-sensitive peptidyl-tRNA forms of the proteins. The dashed arrow indicates the ribosome-associated forms and the solid arrow indicates soluble full length protein forms.

